# NK cells force cytomegalovirus to use hematopoietic cells and immune evasion for dissemination after mucosal infection

**DOI:** 10.1101/618132

**Authors:** Shunchuan Zhang, Finn Grey, Christopher M. Snyder

**Affiliations:** Department of Microbiology and Immunology, Sidney Kimmel Medical College, Sidney Kimmel Cancer Center, Thomas Jefferson University, Philadelphia, PA, 19107; Division of Infection and Immunity, The Roslin Institute, University of Edinburgh, Easter Bush, Midlothian, United Kingdom

## Abstract

Cytomegalovirus (CMV) infects most people in the world and causes clinically important disease in immune compromised and immune immature individuals. How the virus disseminates from the initial site of infection is poorly understood. We used an innovative approach, involving insertion of target sites for the haematopoietic specific miRNA miR-142-3p into an essential viral gene in murine cytomegalovirus. This virus was unable to disseminate to the salivary gland following intranasal infection, demonstrating a strict need for hematopoietic cells for dissemination from the natural site of infection. Viral immune evasion genes that modulate MHC-I expression and NKG2D activation were also required in this setting, as MCMV lacking these genes exhibited impaired dissemination of the viral genome to the salivary gland, and there was no detectable viral replication in the salivary gland. Depletion of T cells rescued the replication of this evasion-deficient virus in the salivary gland. Surprisingly however, the early dissemination to the salivary gland of this evasion-deficient virus, could be rescued by depletion of NK cells, but not T cells. These data are the first to show a profound loss of MCMV fitness in the absence of its MHC-I evasion genes and suggest that they protect the virus from NK cells during hematopoietic dissemination to the salivary gland, where they continued to need the three evasion genes to avoid T cell responses. Remarkably, we found that depletion of NK cells also freed the virus from the need to infect hematopoietic cells in order to reach the salivary gland. Thus, our data show that MCMV adapts to NK cell pressure after intranasal infection by using hematopoietic cells for dissemination while immune evasion genes protect the virus from NK cells during dissemination and from T cells within mucosal tissues.

## Introduction

Cytomegalovirus (CMV), is the most common infectious cause of birth defects in the developed world, leading to hearing loss, vision impairment and cognitive/motor deficits and is estimated to affect 0.5% to 5% of children globally [1-3]. The greatest risk for the most devastating outcomes of congenital CMV infection occur when a pregnant mother experiences a primary infection and the virus disseminates from the site of entry (most likely the oral/nasal cavity) to the fetus [2, 3]. However, infection of the fetus in this circumstance is not universal. In fact, only ∼40% of primary infections during pregnancy result in congenital infection, although the reasons for these drastically different outcomes are unknown. Thus, an understanding of the host/pathogen relationship that governs viral dissemination from the site of entry is critical for the development of effective anti-viral strategies and vaccines.

Primary CMV infection in immune-competent hosts is usually clinically silent, which makes early natural infection difficult to detect and study. Several excellent animal models have been described for investigating CMV infections, including murine (M)CMV, which has been extensively used for *in vivo* studies due to the wealth of available tools. Recent work has identified the nasal mucosa of mice as a natural site of primary MCMV infection [4]. After entry, MCMV must disseminate to the salivary gland, which is a key site of viral persistence and shedding for transmission to new hosts. Thus, understanding the host/pathogen interactions surrounding MCMV dissemination from the nasal mucosa to the salivary gland should provide key information about natural CMV dissemination after primary infection. However, most studies with MCMV, as well as other animal models of CMV infection, have utilized non-physiological routes of infection including intraperitoneal (i.p.), intravenous (i.v.) or subcutaneous (s.c.) inoculation (e.g. [5-7]). After inoculation via the i.p., i.v., and even foot-pad (f.p.) routes, MCMV can directly infect cells in the spleen [5, 6, 8], indicating a direct hematogenous dissemination from the site of inoculation. In contrast, intranasal (i.n.) inoculation, like natural infection, involves no break of the barrier tissue, forcing the virus to go through at least one round of infection before it can disseminate.

Extensive work has suggested that infection of monocytes is important for MCMV dissemination to the salivary gland after foot-pad inoculation [9-12]. Moreover, very recent data has suggested that dendritic cells are important for viral dissemination to the salivary gland after i.n. infection [13], thus supporting a central role for hematopoietic cells in viral dissemination. If MCMV must infect hematopoietic cells for dissemination after i.n. inoculation, we reasoned that evasion of T cell and NK cell responses should facilitate viral spread. All CMVs encode several genes to block the MHC-I antigen presentation pathway. For MCMV there are 3 known genes (m04, m06 and m152), whose protein products act together to interfere with the trafficking of mature MHC-I to the cell surface, consequently protecting infected cells from killing by CD8^+^ T cells [14-16]. Moreover, the m04 and m152 gene products also inhibit NK cell responses [17-20]. However, despite clear *in vitro* evidence for the effectiveness of m04, m06 and m152 [14, 16, 21-24], deletion of all three MHC-I evasion genes had only modest effects on viral dissemination and viral loads *in vivo* after i.p., i.v., or f.p infection [25-28]. In fact, these genes were only found to be critical in three published cases. For MCMV, the Oxenius lab reported that loss of m04, m06 and m152 impaired viral replication in the salivary gland after i.v. infection when mice lacked CD4^+^ T cells, and thus depended exclusively on CD8^+^ T cells for viral control [29]. In addition, the Reddehase group reported that MHC-I evasion genes enhanced the latent MCMV load and played a vital role in viral reactivation from latency in lung explants [30]. Finally, using the Rhesus (Rh)CMV model, Picker, Früh and colleagues reported that evasion of MHC-I antigen presentation and CD8^+^ T cell responses was critical for experimental superinfection (via the subcutaneous route) of Rhesus macaques that had been previously infected by RhCMV, and thus had robust pre-existing immunity [7].

We investigated the need for infection of hematopoietic cells after i.n. infection using novel recombinant strains of MCMV. Consistent with previous work we found that hematopoeitic cells were critical for MCMV dissemination from the nasal mucosa in C57BL/6 mice. In this setting, MHC-I evasion genes enhanced the amount of virus that reached the salivary gland and protected MCMV from CD8^+^ T cells within the nasal mucosa and salivary gland. Surprisingly, our data show that NK cells enforced this requirement: depletion of NK cells or use of BALB/c mice, allowed viral dissemination that did not require infection of hematopoietic cells or MHC-I evasion genes. These data show for the first time that MHC-I modulating genes can be critical for viral dissemination from a natural site of entry to a key site of shedding, providing an explanation for the presence of these genes in the CMV genome. Moreover, our data suggest that early immune responses and the genetic background of the mice directly affect the efficiency of MCMV dissemination to the salivary gland after infection of the nasal mucosa.

## Results

### MCMV must infect hematopoietic cells for dissemination in C57BL/6 mice after i.n. inoculation

MCMV has been proposed to disseminate from the nasal mucosa and other tissues in hematopoietic cells [13, 31]. To test whether infection of hematopoietic cells is necessary after intranasal infection, we constructed a recombinant MCMV containing four repeated targeting sites for the microRNA miR-142-3p in the 3’ untranslated region of the essential viral gene IE3 (MCMV-IE3-142, Figure 1A). As miR-142-3p is exclusively expressed in hematopoietic cells, we predicted that MCMV-IE3-142 would fail to replicate in hematopoietic cells due to targeting of IE3 expression by miR-142, but would replicate to wild type levels in all other cell types. As a control, a second virus was produced containing shuttle vector sequences, but no miR-target sites (MCMV-IE3-015). While both viruses replicated equally well in 3T3 fibroblast cells, only the control virus replicated in macrophages (Figure 1B). To directly visualize the regulation of gene expression by miR-142-3p, a second set of viruses was produced containing GFP in the IE2 locus and either 4 miR-142-3p targeting sites (MCMV-GFP-142) or control vector sequences in the 3’ untranslated region (MCMV-GFP-015) (Figure 1A). Both viruses expressed GFP in infected fibroblast cells, but only the control virus expressed GFP in IC-21 macrophages (Figure 1C) or primary bone marrow macrophages (data not shown). Collectively, these *in vitro* data showed that miR-142 targeting sites severely limited expression of the targeted viral genes and that targeting the essential IE3 gene by miR-142-3p prevented viral replication in hematopoietic cells.

**Figure 1.**
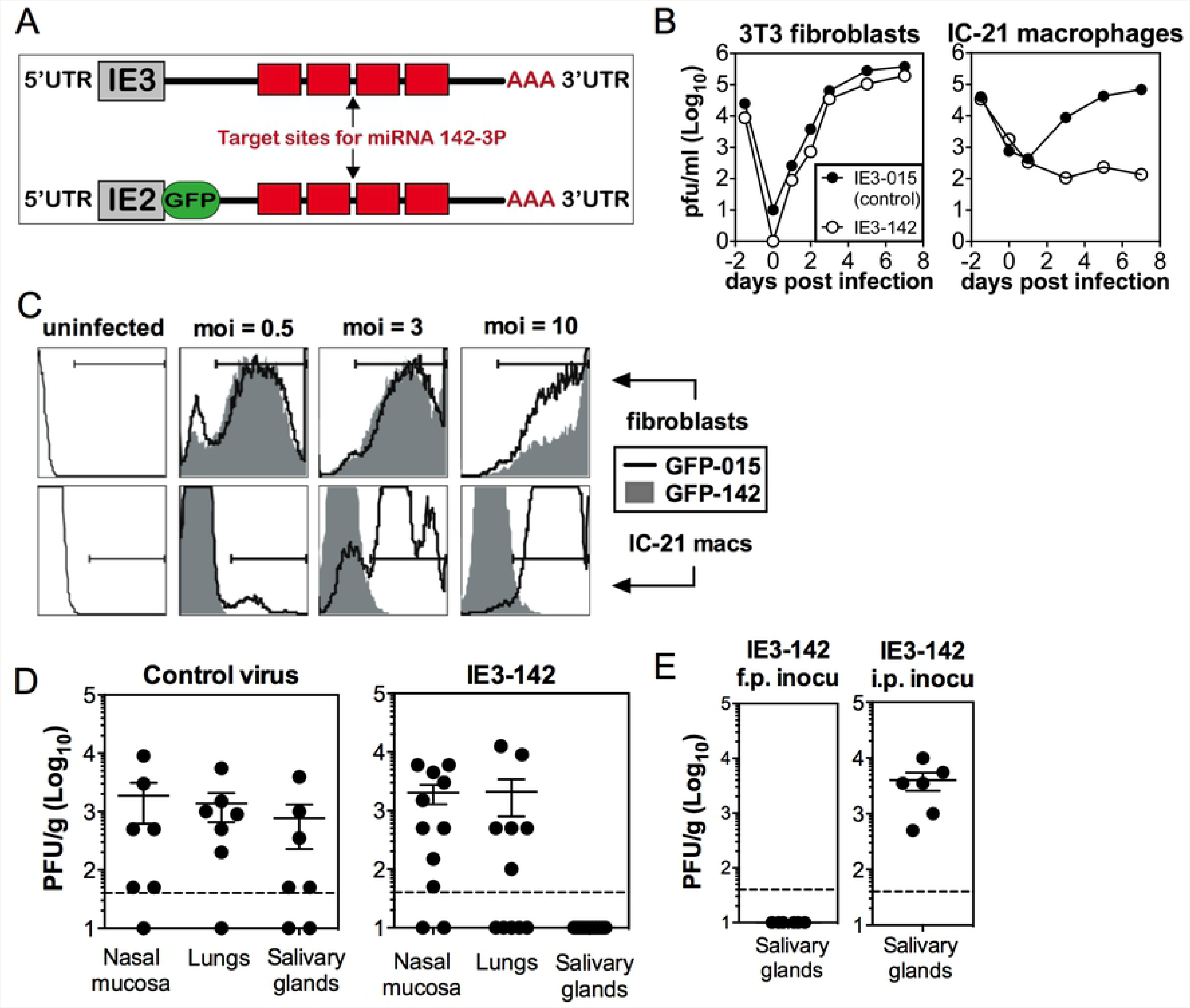
MCMV must use infected hematopoietic cells for dissemination in C57BL/6 mice after i.n. inoculation. **A.** Schematic of miR-142-3p targeted viruses (MCMV-IE3-142 and MCMV-GFP-142). Four target sites for miR-142-3p were inserted into the 3’ untranslated region of the essential viral gene IE3 or into a GFP-SIINFEKL fusion construct inserted into the IE2 locus. Control viruses contain shuttle vector sequences without miR-142-3p target sites in the same location. **B.** Targeting IE3 with miR-142-3p binding sites prevents viral replication in miR-142-3p-expressing macrophages. Multi-step growth curves of the MCMV-IE3-015 control virus and the MCMV-IE3-142 virus in 3T3 fibroblasts and IC-21 macrophages. **C.** Targeting IE3 with miR-142-3p binding sites inhibits gene expression. Regulation of gene expression by miR-142-3p is visualized by GFP expression after infection of 3T3 fibroblasts and IC-21 macrophages with either MCMV-GFP-015 or MCMV-GFP-142 viruses. **D-E.** Productive infection of hematopoietic cells is necessary for viral dissemination after i.n. and f.p. inoculation, but not after i.p. inoculation. Virus titers in the nasal mucosa, lungs and salivary glands at 14 days after i.n. inoculation (**D**) or f.p. or i.p. inoculation (**E**), with 10^6^ MCMV-IE3-142, or control virus MCMV-IE3-015. Each symbol represents an individual animal. The solid line shows the mean titer, and error bars represent the SEM. Dashed lines show the detection limit (50 PFU/g). Data are combined from two independent experiments.

Recent work has shown that the nasal mucosa is a natural site of MCMV entry [4]. Thus, we infected C57BL/6 mice by the intranasal route with either MCMV-IE3-142 or control viruses (either parental wild-type BAC MCMV, or MCMV-IE3-015). All three viruses replicated at the sites of entry (nasal mucosa and lungs). However, two weeks after infection, only control viruses were found to be replicating in the SG (Figure 1D). Similar results were obtained after footpad inoculation, which is considered reflective of infection via licking skin abrasions or biting, another possible natural route of infection for MCMV (Figure 1E). In contrast, infection by the i.p. route, which allows hematogenous spread of cell-free virus [32], enabled MCMV-IE3-142 to replicate robustly in the SG (Figure 1E). Taken together, these data show that MCMV must utilize infected hematopoietic cells to efficiently spread to SG after intranasal infection

### Evasion of MHC-I antigen-presentation and CD8^+^ T cells is critical for viral persistence at sites of entry and replication in the salivary gland

Since MCMV had to infect hematopoietic cells to spread to SG after i.n. inoculation, we considered whether these infected hematopoietic cells must evade immune control in order to disseminate the virus. MCMV encodes three MHC-I evasion genes (m04, m06 and m152), which collectively interfere with the trafficking of mature MHC-I to the cell surface and protect infected cells from killing by CD8^+^ T cells *in vitro* [14, 16, 21-24]. However, despite clear *in vitro* evidence for the function of these MHC-I evasion genes [14, 16, 21-24], previous studies on the *in vivo* impact of losing all three MHC-I evasion genes have revealed only subtle defects in the kinetics of viral replication and clearance, the size and kinetics of the virus-specific CD8^+^ T cell response, or the total latent viral loads after i.p. or f.p infection [25, 26, 28, 33]. Only MCMV’s replication in the salivary glands of CD4^+^ T cell deficient mice and its reactivation from latency in explant cultures have been reported to require these MHC-I evasion gene [29, 30]. To test whether MHC-I immune evasion genes play an important role after infection of the nasal mucosa, a natural site of entry, C57BL/6 mice were infected i.n. with either wild-type MCMV (WT-MCMV) or “Triple Knockout” MCMV (TKO-MCMV) lacking all three MHC-I evasion genes. While WT-MCMV replicated at the entry sites (nasal mucosa and lungs) from 7 days after infection until the end of the experiment at day 28, and also spread to SG within 14 days (Figure 2A), TKO-MCMV was controlled at the sites of entry and failed to spread to SG except in one mouse (Figure 2B). Importantly, depletion of CD8^+^ T cells prior to infection enabled TKO-MCMV to replicate normally and persist for at least 28 days in the nasal mucosa and lungs while also spreading to SG with similar kinetics as WT-MCMV (Figure 2B). These data clearly show that viral evasion of MHC-I antigen presentation and CD8^+^ T cells is crucial for MCMV to effectively spread to SG from a natural site of infection.

**Figure 2.**
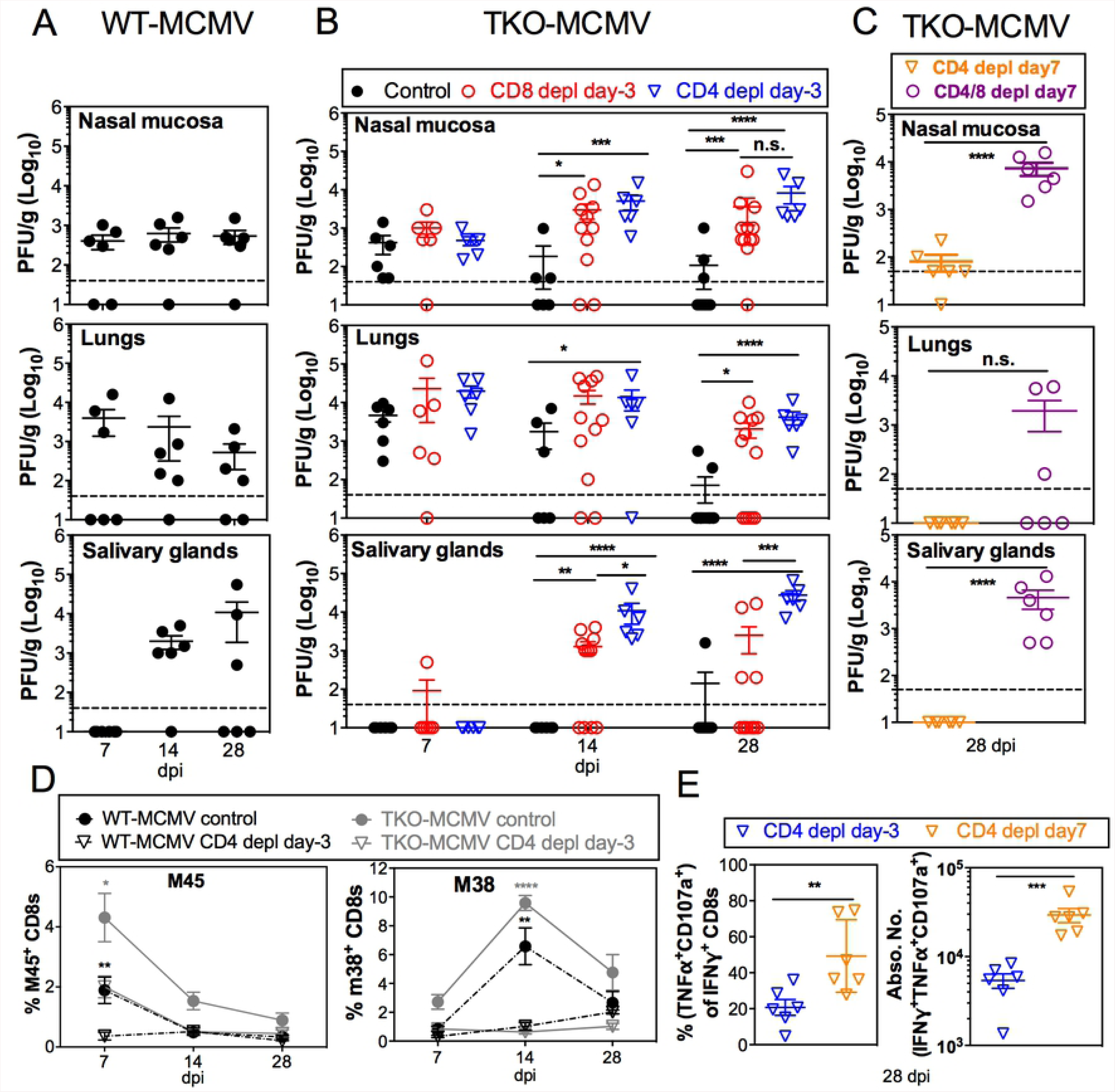
Evasion of MHC-I antigen-presentation and CD8^+^ T cells is critical for viral persistence at site of entry and replication in the salivary gland. **A.** Wild-type MCMV persists in the nasal mucosa and spreads to the salivary gland within 14 days after i.n. infection. Virus titers in the nasal mucosa, lungs and salivary glands of C57BL/6 mice at 7, 14 and 28 days following i.n. inoculation of WT-MCMV. Each symbol represents an individual animal. The solid line shows the mean titer, and error bars represent the SEM. Dashed lines show the detection limit (50 PFU/g). Data are combined from two independent experiments. **B.** Lack of MHC-I evasion genes prevents viral persistence in the nasal mucosa and spread to the salivary gland. Viral titers in the nasal mucosa, lungs and salivary gland 7, 14 and 28 days after i.n. inoculation of C57BL/6 mice infected with TKO-MCMV. Infected mice were treated with an isotype control antibody or depleted of CD4^+^ or CD8^+^ T cells before infection. Data are displayed as in **A** and are combined from two independent experiments. **C.** CD8^+^ T cells can control TKO-MCMV if they are primed in the presence of CD4^+^ T cell help. Virus titers in the indicated organs at day 28 post infection are shown. C57BL/6 mice were depleted of either CD4^+^ T cells or both CD4^+^ and CD8^+^ T cells, beginning at day 7 after i.n. infection of TKO-MCMV. Data are displayed as in **A** and are combined from two independent experiments. **D.** CD8^+^ T cells are reduced in frequency after i.n. infection in the absence of CD4^+^ T cell help. Shown is the frequency of viral tetramer-specific CD8^+^ T cells in the blood of recipients at the indicated time points with or without CD4^+^ T cell depletion before infection. Data show the average frequency of T cells at day 7 (n=9-12), day 14 (n=6-9) and day 28 (n=3-6) after i.n. infection and are derived from one representative experiment of at least 3 independent experiments. **E.** CD8^+^ T cell function is impaired in the absence of CD4^+^ T cell help after i.n. infection, but improved by delaying CD4^+^ T cell depletion until day 7. Each symbol represents an individual animal. The solid line shows the mean value, and error bars represent the SEM. Data are combined from two independent experiments.

Surprisingly, depletion of CD4^+^ T cells also reversed the restriction on viral dissemination, enabling TKO-MCMV to replicate persistently at the entry sites and spread to SG (Figure 2B). In fact, the virus replicated to even higher titers in the salivary gland in CD4^+^ T cell-depleted mice (Figure 2B). CD4^+^ T cells are well-known to play a direct role in the control of MCMV in the salivary gland [34-36] and a recent report has suggested that MCMV must evade CD4^+^ T cells via the viral gene M78 for efficient dissemination to the salivary gland [37]. However, since none of the three viral genes missing in TKO-MCMV (m04, m06 and m152) are known to contribute to evasion of MHC-II or CD4^+^ T cells, we wondered whether CD4^+^ T cell help was needed to develop functional CD8^+^ T cell responses after i.n. infection, rather than to directly control TKO-MCMV. This would be unexpected since previous work suggested that CD4^+^ T cell help plays only a modest role in supporting CD8^+^ T cells after i.p. infection, primarily affecting T cell recall capacity and memory inflation [38, 39]. To test this, we depleted CD4^+^ T cells beginning at day 7 after infection, which should allow CD8^+^ T cells to be primed in the presence of CD4^+^ T cell help. At this time-point, similar titers of WT-MCMV and TKO-MCMV were present in the nasal mucosa and lungs, but neither virus was replicating in the salivary gland (Figure 2A and 2B). This delayed depletion of CD4^+^ T cells enabled control of TKO-MCMV in the nasal mucosa and lungs and prevented any virus from being detected in the salivary gland (Figure 2C). Control was not due to the effects of CD4^+^ T cells in the first week of infection because double depletion of CD4^+^ T cells and CD8^+^ T cells, both beginning on day 7 after infection, restored TKO-MCMV replication in all tissues (Figure 2C). Thus, delayed depletion of CD4^+^ T cells restored control of TKO-MCMV in a CD8^+^ T cell-dependent manner.

Consistent with a role for CD4^+^ T cell help for CD8^+^ T cells, CD4^+^ T cell depletion prior to i.n. infection significantly reduced the frequency and number of MCMV-specific CD8^+^ T cells in the blood (Figure 2D & Figure S1A). Representative gating strategies for these and subsequent data is shown in Figure S2. For precise quantitation of CD8^+^ T cell function per cell we used OT-I T cells stimulated by i.n. infection with MCMV-Ova. Depletion of CD4^+^ T cells prior to infection resulted in impaired cytokine production and degranulation of OT-Is (Figure S1B). In contrast, delaying depletion of CD4^+^ T cells until day 7 after infection significantly increased the frequency and number of functional CD8^+^ T cells (Figure 2E). Thus, CD4^+^ T cell help was critical for the development of functional CD8^+^ T cell responses after i.n. inoculation, which were able to completely prevent TKO-MCMV from replicating in the salivary gland.

### Early dissemination of TKO-MCMV is restored by NK cell depletion

It was possible that MHC-I evasion genes protected MCMV during dissemination to the SG or after it arrived. If MHC-I evasion genes were required during dissemination, we reasoned that we would detect reduced quantities of TKO-MCMV DNA in the SG, which should be rescued by T cell depletion. To specifically assess viral dissemination rather than replication after dissemination, we assessed viral DNA load in the SG 4 days after i.n. inoculation, a time point at which virus-specific CD8^+^ T cells can be detected in draining lymph nodes, but not yet in the SG (Figure 3A). Indeed, the TKO-MCMV DNA load was approximately 10-fold reduced compared to WT-MCMV in unmanipulated C57BL/6 mice (Figure 3B). However, when CD8^+^ T cells, CD4^+^ T cells or both CD4^+^ and CD8^+^ T cells were depleted, TKO DNA load was only marginally increased (about 2-fold) and this did not reach significance (Figure 3B). Moreover, TKO-MCMV DNA was present in similar amounts in draining lymph nodes (mandibular LNs, deep cervical LNs and mediastinal LNs [40, 41]) with or without CD8^+^ T cells (Figure 3C). These data show that dissemination of TKO-MCMV is markedly impaired, but suggest that T cells are not principally responsible, despite their effects on TKO-MCMV persistence in the nasal mucosa and lungs, and replication in the salivary gland (Figure 2). However, depletion of NK cells alone or NK cells and T cells from C57BL/6 mice resulted in complete restoration of early TKO-MCMV dissemination to the salivary gland (Figure 3B). Moreover, mice deficient in either perforin or IFN-γ completely failed to restrict TKO-MCMV dissemination (Figure 3D). Thus, NK cells strongly limited the dissemination of TKO-MCMV through a mechanism that required both perforin and IFN-γ. In contrast, depletion of NK cells did not rescue MCMV replication in the salivary gland by day 14 (Figure 3E), unlike depletion of T cells (Figure 2B). Together, these data show that the MHC-I/NKG2D evasion genes are required to escape NK cell responses during dissemination in hematopoietic cells and to escape T cell responses during replication in the salivary gland.

**Figure 3.**
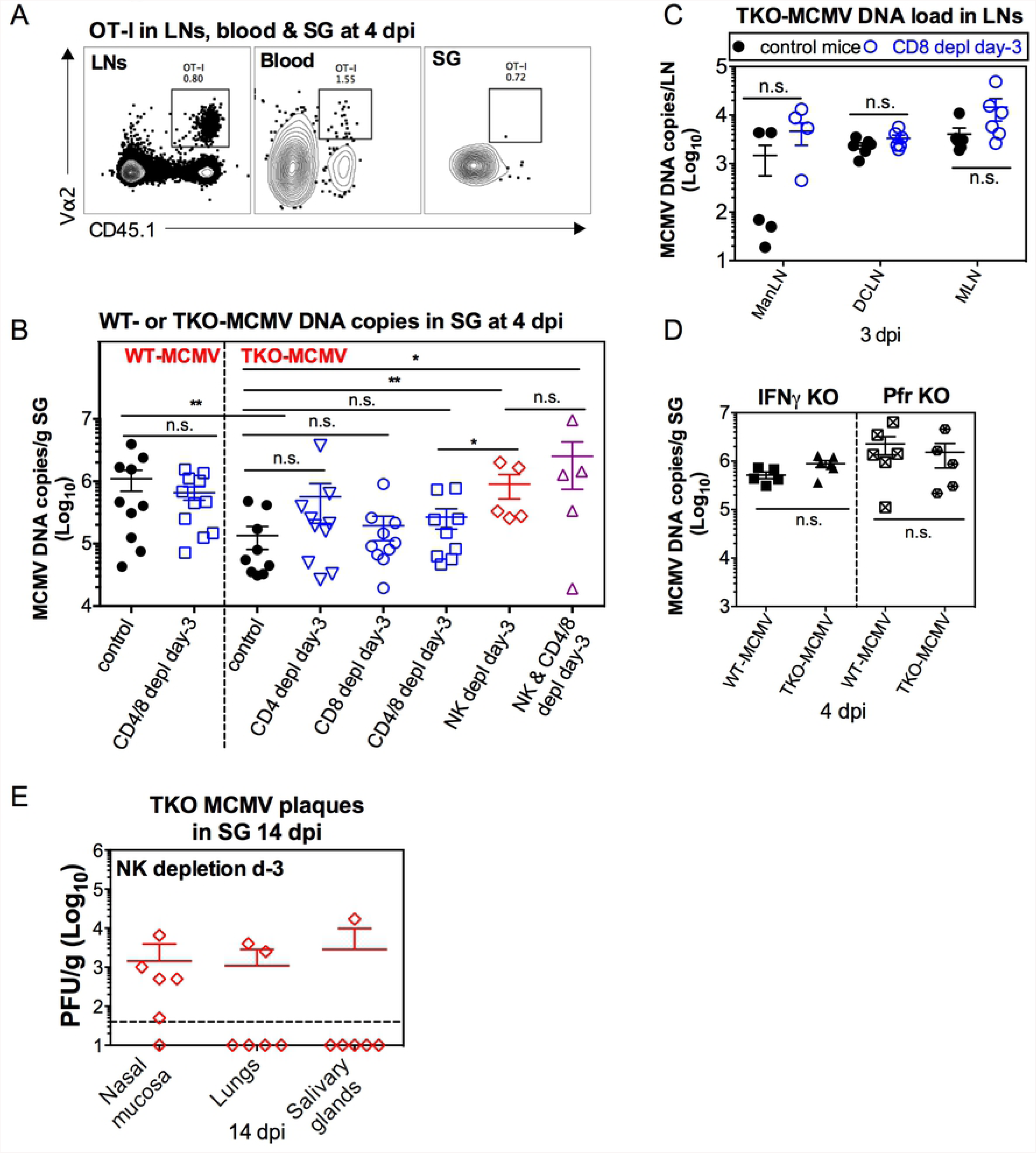
Early dissemination of TKO-MCMV is impaired by NK cells. **A.** MCMV-specific T cells are not present in the salivary gland by day 4 after i.n. infection. Representative FACS plots show OT-I cells in the blood, draining LNs (ManLNs, DCLNs and MLNs) and SG 4 days after i.n. infection with MCMV-OVA. Data show cells in one representative mouse from one experiment. **B.** NK cells, but not T cells, prevent dissemination of TKO-MCMV to the salivary gland after i.n. infection of C57BL/6 mice. C57BL/6 mice were depleted of the indicated cells before i.n. inoculation with TKO-MCMV. Shown are viral DNA copies in the SG 4 days after infection. Each symbol represents an individual animal. The solid line shows the mean value, and error bars represent the SEM. Data are combined from two independent experiments for each condition. **C.** CD8^+^ T cell depletion does not change the TKO-MCMV DNA loads in draining LNs (ManLNs, DCLNs and MLNs). CD8^+^ T cells were depleted or not from C57BL/6 mice. Shown are viral DNA copies in the indicated lymph nodes at 3 days post infection. Data are displayed as in **B** and are combined from 2 independent experiments. **D.** IFN-γ and perforin are essential to prevent dissemination of TKO-MCMV. Shown are copies of viral DNA in the SG following i.n. infection with either TKO-MCMV or WT-MCMV of mice lacking IFN-γ or perforin. Data are displayed as in **B** and are combined from 2 independent experiments. **E.** NK cell depletion before infection does not rescue TKO-MCMV replication in the salivary glands after i.n. infection of C57BL/6 mice. Shown are virus titers in the nasal mucosa, lungs and salivary glands at 14 days post infection. Data are displayed as in **B** and are from one experiment.

### NK cells and T cells enforce the need for hematopoietic cell dissemination

Thus far, our data indicate that MCMV dissemination after i.n. inoculation required infection of both hematopoietic cells and the MHC-I evasion genes in C57BL/6 mice. However, depletion of NK cells and not T cells rescued dissemination of TKO MCMV to the salivary gland. C57BL/6 mice are resistant to MCMV infection as a result of activation of Ly49H^+^ NK cells by viral m157 [42, 43]. Therefore, we wondered whether TKO-MCMV would spread more efficiently in BALB/c mice, which lack Ly49H-expression and robust NK cell responses. Indeed, TKO-MCMV readily spread to the SG in BALB/c mice by day 14 (Figure 4A). Even more remarkably, MCMV-IE3-142 virus was also able to spread to the SG in BALB/c mice and replicate there within 14 days of infection (Figure 4B). Thus MCMV did not need to infect hematopoietic cells or express MHC-I evasion genes to reach the salivary gland in BALB/c mice. If this effect was mediated by the NK cell response, we should be able to restore MCMV-IE3-142 spread to the SG in C57BL/6 mice simply by depleting NK cells. Indeed, depletion of NK cells from C57BL/6 mice prior to i.n. infection allowed MCMV-IE3-142 to reach the SG in nearly all (11 of 12) mice (Figure 4C). Moreover, it accelerated viral dissemination and led to increased copies of MCMV-IE3-142 DNA in the SG as early as day 4 after infection, with no significant difference in viral DNA loads in the SG between BALB/c mice and C57BL/6 mice lacking NK cells (Figure 4D). Thus, the requirement for dissemination within infected hematopoietic cells was dictated by the NK cell response in C57BL/6 mice. Interestingly, depletion of CD4^+^ and CD8^+^ T cells from C57BL/6 mice also partially rescued MCMV-IE3-142 dissemination, allowing salivary gland replication in 5 of 12 mice (Figure 4C), but did not result in increased viral DNA loads by day 4 after infection (Figure 4D). Thus, in the absence of T cell responses MCMV-IE3-142 could eventually reach the SG without using hematopoietic cells in some mice, but the early dissemination was unaffected. Together, these data suggest that early control of MCMV by NK cells forced MCMV to infect and utilize hematopoietic cells for dissemination and that MHC-I/NKG2D evasion genes were required to avoid early NK cell control during dissemination and to productively infect the SG after arrival.

**Figure 4.**
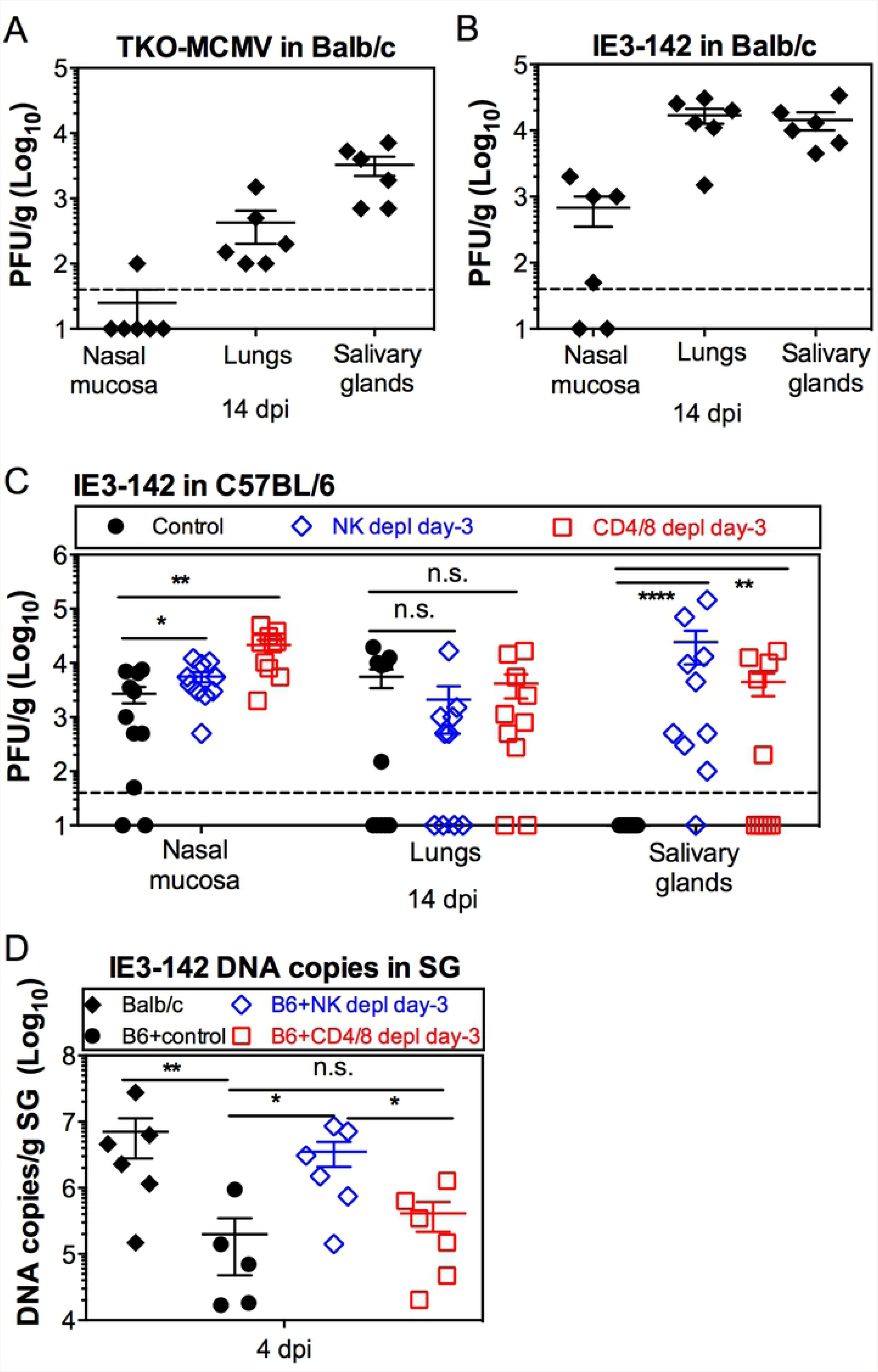
NK cells and T cells enforce the need for hematopoietic cell dissemination. **A.** MCMV does not require MHC-I evasion to reach the salivary gland after i.n. infection of BALB/c mice. Virus titers in the nasal mucosa, lungs and SG of BALB/c mice at 14 dpi after i.n. inoculation of TKO-MCMV. Each symbol represents an individual animal. The solid line shows the mean value, and error bars represent the SEM. Dashed lines show the detection limit (50 PFU/g). Data are combined from at least two independent experiments. **B.** MCMV does not require infection of hematopoietic cells to reach the salivary gland after i.n. infection of BALB/c mice. Shown are virus titers in the nasal mucosa, lungs and SG of BALB/c mice 14 days after i.n. inoculation of MCMV-IE3-142. Data are displayed as in **A**. **C.** Depletion of NK cells or T cells from C57BL/6 mice enables viral dissemination from the nasal mucosa to the salivary gland without infection of hematopoietic cells. C57BL/6 mice were depleted of NK cells or CD4^+^ T cells and CD8^+^ T cells before i.n. infection with MCMV-IE3-142. Shown are the viral titers in the nasal mucosa, lungs and SG 14 days after infection. Data are displayed as in **A**. **D.** Early viral dissemination is affected by NK cell responses. BALB/c mice or C57BL/6 mice with or without depletion of either NK cells or CD4^+^ and CD8^+^ T cells were i.n. inoculated with MCMV-IE3-142. Shown are viral DNA copies in the SG at 4 dpi after infection.

## Discussion

Over the last 6 decades, nearly all work with MCMV, as well as other animal models of CMV infection, have utilized non-physiological routes of inoculation including i.p., i.v., s.c. and f.p., with a few studies utilizing an i.n. route to study mucosal infection. However, recent work directly showed that natural MCMV transmission from infected mothers to pups occurs through the nasal mucosa [4]. Moreover, human CMV is thought to infect via an oral/nasal route and it has been proposed that the nasal mucosa may be the dominant site of entry [4]. Very little is known about the immune response or viral dissemination after a nasal infection and our data reveal a surprisingly complex and previously unappreciated host/pathogen relationship after intranasal inoculation of mice with MCMV. First, and most surprisingly, we found that strong NK cell responses during primary viral infection forced MCMV to use infected hematopoietic cells for dissemination from the entry sites (nasal mucosa and lungs). A lack of NK cells or infection of BALB/c mice enabled non-hematopoietic viral dissemination. Second, our data suggest that MHC-I/NKG2D evasion genes were critical to enable efficient dissemination to the SG and that the impaired dissemination could be restored by depletion of NK cells. Third, we found that MHC-I/NKG2D evasion was required for MCMV to evade T cell responses in order to persist in the nasal mucosa and replicate in the salivary gland. Finally, our data show that CD4^+^ T cell help was required to produce functional and protective CD8^+^ T cells after such mucosal infection. Overall, these data describe a previously unappreciated host/pathogen relationship that develops after infection by the nasal route.

Our data are the first to describe a vital role for MCMV’s MHC-I evasion genes during primary CMV infection. Previous work in the MCMV model described subtle improvements in viral fitness and latent loads as a result of CD8^+^ T cell evasion, but no severe defects in infection or dissemination for viruses lacking all 3 known MHC-I evasion genes (m04, m06 and m152) [25-28]. Only MCMV’s replication in the salivary glands of CD4^+^ T cell deficient mice and its reactivation from latency in explant cultures have been reported to require these MHC-I evasion genes [29, 30]. However, like most previous studies of MCMV immunobiology, these studies all utilized routes of infection (i.p., i.v. or f.p.) that involve breaking or avoiding a barrier tissue and allow MCMV to directly infect cells in the spleen [5, 6, 8]. This indicates that MCMV likely had direct access to the blood from the site of inoculation. One previous study showed a critical *in vivo* role for evasion of CD8^+^ T cells by RhCMV after subcutaneous infection, but only if animals were previously infected with RhCMV and therefore contained a robust and pre-existing T cell response [7]. In contrast, our data show that primary MCMV infection via a natural mucosal barrier tissue necessitated the expression of these genes in order for the virus to persist in the nasal mucosa, disseminate to the salivary gland and replicate in the salivary gland. As might be predicted, these genes protected MCMV from T cells during replication in the nasal mucosa and salivary gland (Figure 2). In this context, it is interesting to note that recent work identified the viral M78 protein as responsible for reducing MHC-II expression on infected cells, which was required for efficient infection of the salivary gland after intranasal inoculation [37]. Unexpectedly however, dissemination from the nasal mucosa to the salivary gland required the m04, m06 and m152 genes to limit NK cell control of the virus. Previous work has shown that the m04 and m152 genes contribute to evasion of NK cell responses [17-20, 44-46]. The m04 protein complexes with MHC-I and allows increased surface expression of total MHC-I even in cells expressing m152 and m06, to avoid NK cell activation [17]. In addition, the m152 gene down-regulates the RAE-1 ligands for the NKG2D activating receptor, consequently inhibiting NK cell responses [19, 20, 44-46]. Future work will be needed to define the individual roles of viral m04 and/or m152 in improving viral dissemination. Together, these data suggest that MCMV’s MHC-I/NKG2D evasion genes play a vital role in avoiding NK cells during hematopoietic cell-mediated spread from mucosal tissues to the salivary gland, while also protecting replicating MCMV from T cells within mucosal tissues.

We were most surprised to find that, even in the presence of m04, m06 and m152, NK cells forced MCMV to use hematopoietic cells as carriers in C57BL/6 mice. Previous work has described hematopoietic cells, including monocytes, macrophages and dendritic cells, as important carriers for CMV dissemination [4, 9, 13, 47, 48]. Our miR-142-3p targeted strains could not replicate in hematopoietic cells and could not disseminate to the salivary gland in intact C57BL/6 mice (Figure 1). Importantly, this was also true after f.p. infection, but not i.p. infection, indicating that the route of infection dictated the necessity for hematopoietic involvement. These results are consistent with a recent report showing that MCMV used dendritic cells to disseminate to the SG after i.n. inoculation [13] and previous work showing that recruitment of monocytes to the footpad was key for efficient dissemination after f.p. infection [9-12]. However, depletion of NK cells removed the need for hematopoietic cell infection in order for MCMV to disseminate to the salivary gland (Figure 4C). This dramatic effect is likely due to the interaction of host Ly49H (in C57BL/6 mice) and viral m157, as hematopoietic cell infection was not needed in BALB/c mice (Figure 4A). NK cell activating and inhibitory receptor expression varies greatly across genetically diverse populations [49], and other NK cell activating receptors are known to be triggered by MCMV-infected cells (e.g. Ly49P^MA/My^, Ly49P1, Ly49D2, and Ly49L) [50]. Moreover, the specific combination of inhibitory and activating receptors may affect the outcomes as shown by data that the inhibitory Ly49C could inhibit the activation of NK cells through Ly49H [51]. Thus, it is likely that NK cells and viral immune evasion genes play a variable role across an outbred population in regulating the route and pace of viral dissemination after natural infection.

Although our study revealed a critical role for hematopoietic cell infection for dissemination in C57BL/6 mice, it is not clear how MCMV disseminates from the nasal mucosa when NK cells or T cells were depleted or in BALB/c mice. NK cell depletion clearly improved the dissemination efficiency of TKO-MCMV (Figure 3) and MCMV-IE3-142 (Figure 4) in C57BL/6 mice, resulting in increased viral DNA loads in the salivary gland by day 4 after infection. The ability of these viruses to reach the salivary gland after NK cell depletion or in BALB/c mice may imply that it can disseminate by cell-free viremia, or perhaps by infected endothelial cells that are sloughed into the blood stream as they become cytomegalic, a mechanism that was first proposed by Goodpasture and Talbot almost 100 year ago [52]. Future work will be aimed at defining the source of virus that arrives in the salivary gland in each of these settings.

Our data also unexpectedly revealed a critical role for CD4^+^ T cell help after i.n. inoculation. Although CD4^+^ T cell help for CD8^+^ T cells is well established in multiple infectious models (e.g. [53-66]), previous work in the MCMV model had shown that CD4^+^ T cell-deficient mice mounted a remarkably intact CD8^+^ T cell response to MCMV after i.p. or i.v. inoculation [29, 39, 67]. However, after i.n. infection, mice lacking CD4^+^ T cells produced few functional anti-viral CD8^+^ T cells and were unable to limit the dissemination of TKO-MCMV (Figure 2). Previous work has suggested that MCMV-specific CD8^+^ T cells are primed by cross-presentation after i.p. inoculation [68, 69], a route of infection that produces high viral titers in the first few days of infection. In contrast, it is possible that CD8^+^ T cell priming after i.n. infection relies more heavily on direct presentation. Alternatively, cross-presenting dendritic cells in the draining lymph nodes after i.n. MCMV infection may depend on CD4^+^ T cell help for activation and licensing. Future work will be needed to dissect the specific requirements for CD4^+^ T cells after i.n. infection.

Collectively, our data suggest that MCMV is forced by early NK cell responses, and even somewhat by T cell responses, to use the hematopoietic cells for dissemination from the nasal mucosa and lungs. In this setting, the m04, m06 and m152 evasion genes became vital for MCMV to evade NK cell responses to efficiently reach the salivary gland, where they continued to need the three evasion genes to avoid T cell responses and replicate. These data provide the first *in vivo* evidence for a vital role of these immune evasion genes in wild-type, previously uninfected animals and also describe how MCMV can avoid NK cell pressure by utilizing hematopoietic cells to facilitate dissemination.

### Experimental methods

#### Mice

Six to seven-week old mice were used for all experiments. C57BL/6J mice and BALB/c mice were purchased from the Jackson Laboratory and used directly. OT-I transgenic mice (C57BL/6-Tg(TcraTcrb)1100Mjb/J), CD45.1 mice (B6.SJL-Ptprc^a^ Pepc^b^/BoyJ), IFN-γ knock-out mice (B6.129S7-Ifng^tm1Ts^/J) and perforin knock-out mice (C57BL/6-Prf1^tm1Sdz^/J) were purchased from the Jackson Laboratory and maintained in our animal colony. Protocols were approved by the institutional animal care and use committee of Thomas Jefferson University. All mice were housed in a standard pathogen-free animal facility.

#### Viruses

Murine cytomegalovirus strains including the wild-type BAC-derived MCMV strain (MW97.01, called WT-MCMV throughout) [70], TKO-MCMV strain (triple knock out of m04, m06 and m152) [27] and MCMV-Ova (which expresses the cognate SIINFEKL peptide) have been previously described [71, 72]. They were propagated in M2-10B4 cells as previously described [73].

#### Cell culture

M2-10B4 cell line was purchased from ATCC. M2-10B4 cells were cultured in growth media (RPMI-1640 medium with L-glutamine (Mediatech/Cellgro, reference #: 10-040-CV), supplemented with 10% FBS and 100 units/ml Penicillin, 100 µg/ml Streptomycin) at 37°C with 5% carbon dioxide.

#### Experimental infection

Mice under anesthesia were either infected by the i.n. route with 10^6^ PFU MCMV in 20 μL PBS (10 μL per nare), by the i.p. route with 10^6^ PFU MCMV in 100 μL PBS, or by the f.p. route with 10^6^ PFU MCMV in 30 μL PBS. All experiments were approved by the Thomas Jefferson University Institutional Animal Care and Use Committee.

#### Generation of miR-142 cell tropism specific MCMV virus

miR-142 virus and control virus were constructed using the λ-derived linear recombination system in combination with the pSM3fr MCMV bacterial artificial chromosome in the *Escherichia coli* strain DY380 [74]. Sequence containing 4 repeated target sequences with complete complementarity to miR-142-3p were synthesized and inserted 86 bases into the 3’UTR of the IE3 gene (nucleotide coordinate - 177898) of MCMV using linear recombination. Vector sequence containing the FRT flanked kanamycin cassette was inserted into the same region, with the kanamycin cassette removed from both viruses by FLP recombination. miR-142-3p target sequence with complementary sequences highlighted with underline – AGTCGAC<u>TCCATAAAGTAGGAAACACTACA</u>CGAT<u>TCCATAAAGT AGGAAACACTACA</u>ACCGGT<u>TCCATAAAGTAGGAAACACTACA</u>CGAT<u>TCCATA AAGTAGGAAACACTACA</u>ACCGGT. The recombinant viruses have been checked for insertion by restriction analysis, Southern blotting, and sequencing.

#### Multi-step growth curves *in vitro*

3×10^5^ IC-21 macrophages or 2×10^5^ 3T3-fibroblasts were separately seeded into 6-well plates. One day later, cells were infected with either MCMV-IE3-142 or MCMV-IE3-015 at a multiplicity of infection (moi) equalling 0.1 for 2 hours without centrifugal enhancement. After 2 hours, supernatant was collected for input virus titer (labelled day - 1) and cells were washed with fresh media. Cells were scraped from duplicate wells immediately after the wash (day 0) and on days 1, 3, 5 and 7 after infection.

#### Detection of GFP expression by flow cytometry

To visualize the regulation of gene expression by miR-142-3p, MCMV-GFP-142 and MCMV-GFP-015 were used to infect IC-21 macrophages or 3T3-fibroblasts respectively with moi = 0.5, 3 or 10. The following day, cells were fixed and collected to determine GFP expression by flow cytometry.

#### Cell depletions *in vivo*

In some experiments, CD4^+^ T cells, CD8^+^ T cells and/or NK cells were depleted 3 days before infection or 7 days after infection. Depletions were conducted by i.p. injection on days 3, 2 and 1 before infection or 7, 8 and 9 days after infection using 0.2 mg of anti-mouse CD4 mAb (clone GK1.5), anti-mouse CD8β mAb (clone 53-5.8) and/or anti-mouse NK1.1 mAb (clone PK136). Depletions were then maintained for the duration of the experiment by weekly injections with 0.15 mg (GK1.5) or 0.2mg (PK136 and 53-5.8) of antibody. All depleting antibodies were purchased from Bio-x-Cell. Depletions were confirmed by staining for CD4 (clone RM4-4), CD8α (clone 53-6.7) or NKp46 (clone 29A1.4).

#### Virus titration

Nasal mucosa, lungs and salivary glands (SG) were collected at indicated time points post infection and frozen. Nasal mucosa was collected as previously described [40]. Twenty percent homogenates (w/v) were prepared from each collected tissue for virus quantification by plaque assay [40]. Briefly, tissues were weighed and homogenized using a pestle with a small amount of sterile sand in a 1.5 ml centrifuge tube, then suspended with RPMI supplemented with 10% FBS, 100 Units/mL penicillin, and 100 μg/mL streptomycin. Supernatants from the homogenate were collected after centrifugation (2400 xg, 10 min) and viral plaque assay was performed on M2-10B4 cells.

#### Adoptive transfer of OT-I T cells

For adoptive transfer of OT-I T cells we used OT-I transgenic mice expressing CD45.1 as donor cells. Splenocytes containing 5000 OT-I cells from naïve transgenic mice were injected i.v. into sex-matched congenic recipients via the retro-orbital sinus suspended in 100 μl PBS. The following day, recipients were i.n. infected with 10^6^ PFU MCMV-Ova.

#### Lymphocytes isolation, antibodies, tetramer staining, intracellular cytokine stimulation (ICS) and FACS analysis

Lymphocytes from the blood were collected from the retro-orbital sinus and mixed with 10 μl heparin (1000 units/ml). Lymphocytes from the spleen were collected by homogenization of the spleen through a 70 μM filter and suspended in T cell medium (RPMI-1640 medium with L-glutamine, 10% FBS, 100 units/ml Penicillin, 100 μg/ml Streptomycin, and 5 x 10^-5^M β-mercaptoethanol). CD4^+^ and CD8^+^ T cells were identified by antibodies specific for CD3 (clone 17A2), TCRβ (clone H57-597), CD4 (clone RM4-4) and CD8α (clone 53-6.7). OT-I donor cells were further distinguished from recipient cells by CD45.1 (clone A20) and TCR Vα2 (clone B20.1). MHC-I-tetramers loaded with peptides from M45 and M38 were generated at the NIH tetramer core facility (http://tetramer.yerkes.emory.edu/) and used to identify MCMV-specific CD8^+^ T cells as described previously [75]. For assessment of cytokine production after stimulation, splenocytes were stimulated with 1 μg/ml M38_316-323_ peptide (Genemed Synthesis, Inc), 1 μg/ml Brefeldin A (GolgiPlug, BD, Bioscience) in the presence of antibody specific for CD107a (clone 1D4B) at 37°C for 3 hours. Cells were chilled on ice, and live cells were discriminated with Zombie Aqua (Biolegend) prior to staining for expression of CD3 (clone 17A2), and CD8α (clone 53-6.7). Finally, splenocytes were fixed and permeabilized with Cytofix/Cytoperm (BD Biosciences), following the manufacturer’s instructions, and stained for intracellular TNF-α (clone MP6-XT22) and IFN-γ (clone XMG1.2). All antibodies were purchased from Biolegend and cells were collected on BD Fortessa and analyzed with FlowJo sofeware (TreeStar).

#### DNA extraction and quantitative real-time PCR (qPCR)

For extracting DNA from the SG, 50 μl from a twenty percent homogenate (w/v) was used. For mandibular lymph nodes (manLNs), deep cervical lymph nodes (DCLNs) and mediastinal lymph nodes (MLNs), the whole lymph node was used after homogenization in RPMI-1640 medium using a pair of needles. In both cases, DNA was extracted from the whole lymph node and SG using the Puregene core kit A (Qiagen) and following the manufacturer’s instructions for extraction of DNA from tissues. RNA was removed by adding RNase A solution and DNA was eluted with 30 μl distilled water. Extracted DNA was quantified by nanodrop and two microliters DNA were used as a template in each qPCR reaction. The qPCR targeting MCMV-E1 gene was performed as previously described [68]. The genome copy numbers were calculated based on a standard curve of a plasmid containing the MCMV-E1 gene.

#### Statistical analysis

Data in all experiments is pooled from at least two independent experiments. Error bars represent the standard error of the mean (SEM) unless specified otherwise in the figure legend. A two-tailed Student’s t test was used for statistical analysis for pairwise comparisons. All data analyses were performed in Graphpad Prism 6. For all statistical analysis, *p < 0.05, **p <0.01, ***p <0.005.

## Acknowledgments

This work was supported by grant AI106810 awarded to C.M.S.

## Author contributions

S.C.Z., C.M.S. and F.G. conceived of the study. S.C.Z. and C.M.S. performed the experiments. F.G. designed and constructed the MCMV-IE3-142 and MCMV-GFP-142 as well as control viruses, and confirmed miR-1423p expression in macrophages. All authors contributed to the manuscript preparation.

